# CD38^hi^CD19^dim^ cells in lymph nodes predict favorable prognosis in patients with stage III melanoma receiving adjuvant PD-1-blockade

**DOI:** 10.1101/2025.11.03.685725

**Authors:** Ieva Ailte, Sudhir Kumar Chauhan, Lina Prasmickaite, Assia V. Bassarova, Amanda Poissonnier, Marike Feenstra, Marta Nyakas, Bjarne Johannessen, Truls Ryder, Robert Hermann, Lars Frich, Arild Holth, Else Marit Inderberg, Henrik Jespersen, Vivi Ann Flørenes, Jon Amund Kyte, Gunhild M. Mælandsmo, Eivind Valen Egeland

## Abstract

**Background:** Adjuvant immunotherapy has significantly improved survival for patients with stage III cutaneous melanoma, yet a fraction of patients will not benefit from immune checkpoint inhibitors (ICI). The tumor microenvironment plays a pivotal role in generating durable responses to ICI. By analyzing the cellular composition of tumor-associated subsets, key immune components essential for promoting an anti-tumor environment can be pinpointed. This will allow for both patient stratification and identification of biomarkers associated with improved patient outcome.

**Method:** Regional lymph nodes were obtained from patients with stage III melanoma at surgery (n=29). Patients eligible for anti-PD-1 therapy (αPD-1; pembrolizumab or nivolumab) received adjuvant treatment for up to one year. CyTOF was used to determine cellular composition in pre-treatment surgical specimens. Bulk gene expression data generated by NanoString from patients receiving surgery without adjuvant therapy (n=125) was implemented for evaluating trends observed in the CyTOF dataset.

**Results:** Although no significant differences were observed across major hierarchical immune cell types between patients who developed distant metastasis after surgery and those that did not, an increased proportion of CD103^+^PD-1^+^CD8^+^ (T_RM_) T cells and plasmablast-like CD38^hi^CD19^dim^ cells were associated with improved prognosis in the CyTOF cohort. In the untreated cohort, a subset of patients defined as *“Ultra-cold”* (< 2.5 % tumor-infiltrating lymphocyte (TIL) scored by a pathologist) had significantly worse outcome than those with higher TIL infiltration. This *Ultra-cold* TIL group was associated with reduced B cell score, but not CD8^+^ T cell score, as well as reduced expression of activation genes like *CD38*.

**Conclusion:** In this study, CD103^+^PD-1^+^CD8^+^ (T_RM_) T cells and plasmablast-like CD38^hi^CD19^dim^ cell populations were found to be strongly associated with prolonged distant metastasis-free survival in regional lymph nodes from patients with stage III melanoma treated with αPD-1. This suggests an association between progression and infiltration of these cell types at baseline and highlights the potential of using immune cell subsets as prognostic biomarkers.

**What is already known on this topic** – Patients being diagnosed with stage III melanoma will receive immune checkpoint inhibitors but will often be cured by surgery alone. Selection of which patients might benefit from treatment is still unresolved. Accurate biomarkers would aid in treatment stratification, to avoid overtreatment, and unnecessary toxicities.

**What this study adds** – This study highlights how baseline CD103^+^PD-1^+^CD8^+^ (T_RM_) T cells and plasmablast-like CD38^hi^CD19^dim^ cell populations in the regional lymph node, is strongly associated with improved outcome in patients receiving anti-PD-1 therapy.

**How this study might affect research, practice or policy** – The strong association between baseline plasmablast-like cell infiltration in RLN and prolonged distant metastasis free- survival, highlights this cell type as a potential treatment stratification criterion to identify patients with good prognosis.

## Introduction

Melanoma, although accounting for less than 5 % of all skin cancers, represents the most lethal form of skin cancer (1). Early detection of local disease results in a high 5-year survival rate, ranging from 85-100 % (1). Conversely, once melanoma progress from the skin to regional lymph nodes (stage III), the historical risk of relapse and subsequent development of distant metastases (stage IV) is high (2). This progression is associated with a drastically reduced 5-year survival, between 10-30 %.

Immune checkpoint inhibitors (ICI) have revolutionized the treatment of metastatic melanoma, with long-term survival now achievable for more than half of the patients (3). In stage III disease, adjuvant ICI substantially reduces the risk of relapse and distant metastasis, but cancer recurrence is still prevalent and treatment carries a risk of severe or permanent immune-related toxicity (4). Reliable tools to successfully predict benefit of treatment or which patients will develop distant metastasis are therefore urgently needed. Several studies have highlighted the microenvironment as a key for achieving effective anti-tumoral response (5, 6). In particular, activated T cell populations have been emphasized as important drivers of response to ICI, while also distinct cell subtypes/activation states have been associated with response (7–9). Nevertheless, the mechanisms associated with immune escape, treatment resistance and development of distant metastasis are not fully understood. While several studies have tried to identify biomarkers associated with poor prognosis (10–12), few studies have been performed, and no biomarkers have been established for stage III melanoma. By improving our understanding of tumor and microenvironmental changes in the context of ICI treatment and disease progression in stage III melanoma, novel prognostic and predictive biomarkers for response to ICI can be identified.

In the present study, biopsies from stage III melanoma were collected from regional lymph nodes to explore their immune- and tumor cell composition and further evaluate how their functional properties differ between patients developing distant metastasis following anti-PD- 1 treatment (αPD-1) and those not presenting metastatic disease.

## Materials and methods

### Patient cohort

Patients treated for stage III melanoma localized to the regional lymph node (RLN) at Oslo University Hospital (OUS) between 2020-2022 were included in the study cohort. Only patients with lymph node metastasis >1 cm were selected for the study. Eligible patients received standard treatment consisting of surgery followed by αPD-1 therapy, administered as either pembrolizumab or nivolumab for up to one year. For a subset of patients, adjuvant therapy was omitted due to old age, immunosuppressed condition or poor compliance. Biopsies were collected from treatment-naïve patients at the time of surgery.

### Tissue processing

Fresh tissue samples were first disintegrated into small pieces with a scalpel, before being dissociated into single cell solution by enzymatic digestion using DMEM-F12 (Gibco, Life Technologies, Carlsbad, CA, USA) supplemented with 10 % heat-inactivated fetal bovine serum (FBS), 50µg/ml DNase and 1 mg/ml Collagenase IV (all from Sigma-Aldrich, St.Louis, MO, USA) at 37 °C for 1 hour. The suspension was then passed through a 100 µm strainer and diluted in washing medium (DMEM-F12 supplemented with 2 % FBS), followed by centrifugation at 400 *g* for 10 minutes. Samples were cryopreserved in aliquots of 2 × 10^6^ cells resuspended in DMEM-F12 supplemented with 70 % FBS and 10 % DMSO.

### CyTOF staining protocol

Cryopreserved samples were rapidly thawed in a 37°C water bath, then mixed with pre-warmed thawing media (RPMI 1640 (Gibco) supplemented with 10 % FBS). Then cells were centrifuged at 500 *g* for 5 min and resuspended in thawing media supplemented with 100µg/ml DNase, before being placed in an incubator at 37°C for 30min. To identify live cells, cells were stained with Cisplatin in PBS (Gibco) for 5 min at room temperature, followed by incubation in FcR blocking solution (Human TruStain FcX™, BioLegend) to prevent unspecific binding. Extracellular antibodies (Supplementary Table 1) were mixed in the staining buffer containing 0.5 M EDTA (Invitrogen, Life Technologies) in Ca²[/Mg²[- free PBS supplemented with 1 % sterile-filtered bovine serum albumin. Standard staining conditions were 100 µl antibody mix and 100 µl cell suspension containing ∼1[×[10^6^ cells/ml. After incubation with antibodies for 30 minutes at 4°C, cells were washed in 1 mL staining buffer. Then, cells were fixed and permeabilized with eBioscience™ Foxp3/Transcription Factor staining buffer set (Invitrogen), before they were stained with antibodies against intracellular targets (Supplementary Table 1) following manufacturer’s protocol. Finally, cells were incubated with nucleic acid intercalator Iridium (Standard BioTools) for 20 min, washed in Cell Acquisition Solution (CAS; Standard BioTools, South San Francisco, CA, USA) and filtered. Single cell suspensions stained with metal-conjugated antibodies were analyzed by Cytometry by Time-Of-Flight (CyTOF®) on the Helios™ system at the Flow Cytometry Core Facility, Montebello node at OUS. EQ™ Four Element Calibration Beads (Fluidigm) were added at 1:10 dilution immediately before sample injection into CyTOF system.

### CyTOF data analysis

After acquisition, events were normalized, and calibration beads were removed by CyTOF software v7.0 (Standard BioTools). PeacoQC automated quality control to remove abnormal events was performed in CellMass Cytobank software (v6.0). Manual gating of live singlets on PeacoQC-controlled samples was then done based on four Gaussian parameters (center, offset, width, residual) and Cisplatin (Live/Dead) & Iridium (nuclear) stain. The obtained quality controlled single live cell data were analyzed by FlowJo (v10.10.0, Ashland, OR, USA) and CytoBank softwares. Following data normalization and clean-up, the data from all samples were concatenated into a single file for UMAP dimension reduction analysis, before patient samples were re-identified in the concatenated file. Main cell populations were identified by lineage marker expression within main UMAP clusters (Supp. Fig. S1), while sub-populations were gated manually (Supp. Fig. S2).

### CyTOF panel

To identify central cell populations within the RLN tumor-immune microenvironment, a 35- parameter antibody panel for characterization was established in-house. Metal-conjugated antibodies were used as highlighted in (Supp. Table S1), except for antibodies targeting Melan-A and AXL, where carrier-free antibodies were conjugated to metals by Maxpar® X8 Antibody Labeling Kit (Standard BioTools) according to manufacturer’s protocol.

### FlowSOM analysis

FlowSOM (Flow Self-Organizing Map) (13) was used for unsupervised analyses of the CyTOF data. Prior to applying FlowSOM, the melanoma subpopulation was extracted from the concatenated dataset. FlowSOM was applied using default settings and 13 distinct cellular markers expressed in melanoma cells; CD47, AXL, CD80, ARG1, CD44, CD73, Melan-A, CD39, PD-L1, HLA-ABC, CD163, β-catenin and HLA-DR.

### Transcription analysis

Formalin-fixed paraffin-embedded (FFPE) blocks form regional lymph node metastasis from patients with stage III metastatic melanoma who did not receive adjuvant treatment between 2010-2017 were retrieved from the archives at Department of Pathology, OUS, and subjected to NanoString analyses. H&E diagnostic slides were used for selecting areas containing tumor and immune cells for each single patient, before RNA was extracted from the corresponding slides from a total of 125 samples. RNA was generated using the High Pure FFPET RNA isolation kit (Roche, Basel, Switzerland), before the nCounter PanCancer IO 360™ (IO360) panel was applied to all samples. Samples were run on the NanoString nCounter® platform at Experimental Diagnostics, Department of Pathology, OUS.

Quality check was performed using the nSolver Analysis Software (v4.0.70), before counts were processed in R using the approach established in Bhattacharaya et al (14).

### Statistical analysis and figure generation

Statistical analysis and figure generation were performed in R (v4.3.1) programming language with RStudio (2023.09.0+463). Statistical significances were determined using the Wilcoxon non-parametric test using the R package *rstatix*. *P* ≤ 0.05 was considered significant, and labeled with * *P* ≤ 0.05, ** *P* ≤ 0.01, *** *P* ≤ 0.001, **** *P* ≤ 0.0001.

## Results

### Cellular composition of melanoma lymph node biopsies

To characterize melanoma metastasized to the regional lymph node, we collected 29 biopsies from patients with operable stage III disease (Table 1). Initially, UMAP analyses were performed to obtain an unbiased identification of tumor and immune cell composition (Fig. 1a.). This allowed the identification of five major cell clusters, each contributing at least 1 % of the total number of live cells (Fig. 1b). Melanoma cells represented the largest population (27.1 % of all cells), followed by CD8^+^ T cells (24.3 %), CD19^+^ cells (20.8 %), and CD4^+^ T cells (18 %). We also identified minor populations of CD33^+^ myeloid cells (1.7 %), as well as a smaller subset of other CD45^+^ cells (labeled “Other Immune”, ∼1.4 %).

**Figure 1.**
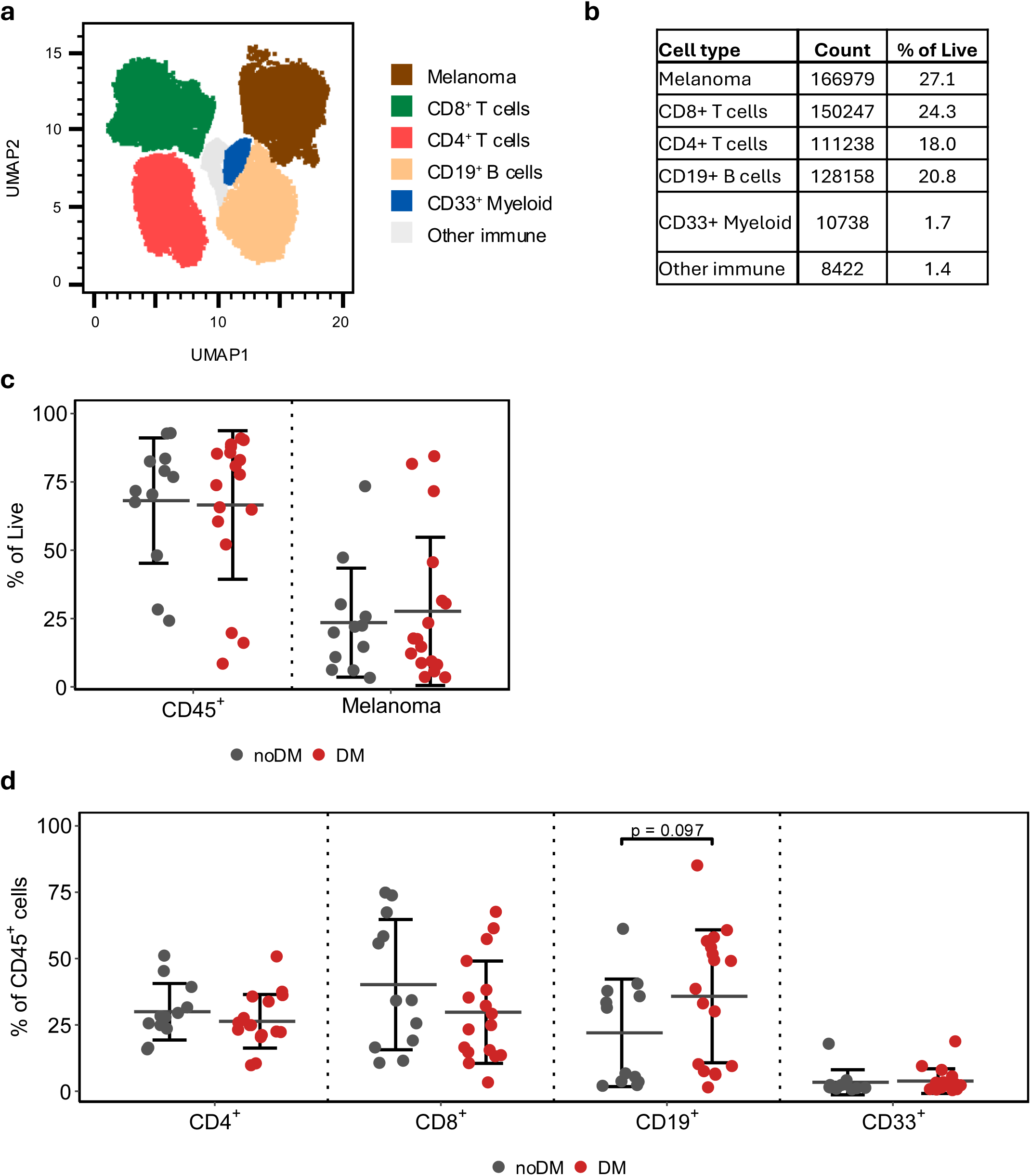
Cellular composition of melanoma lymph node biopsies. **a.** UMAP of single cells from 29 lymph node biopsies from stage III melanoma. Main cell populations are highlighted. **b.** Cell counts and corresponding mean frequencies for main cell populations in a. **c.** The relative proportion of melanoma and CD45^+^ immune cells as a percentage of all live cells in the corresponding biopsy. **d.** The relative proportion of the major immune cell populations across biopsies. Mean±SD as a fraction of CD45^+^ cells. noDM (n = 12); DM (n= 17).

**Table 1.** Clinicopathological parameters and associations with distant metastasis.

|  |  | All patients |  | Distant metastasis |  | No distant metastasis |  | P-value |
| --- | --- | --- | --- | --- | --- | --- | --- | --- |
|  |  | n | % | n | % | n | % |  |
| **Patients** |  | 29 | 100 | 17 | 59 | 12 | 41 |  |
| **Age** | Median (IQR) | 71 (21) |  | 71 (20) |  | 69 (20.5) |  | 0.60<sup>a</sup> |
| **Sex** | Female | 10 | 34 | 4 | 40 | 6 | 60 | 0.24<sup>b</sup> |
|  | Male | 19 | 66 | 13 | 68 | 6 | 32 |  |
| **Melanoma subtype** | Superficial | 8 | 50 | 5 | 63 | 3 | 38 | 0.57<sup>b</sup> |
|  | Nodular | 8 | 50 | 7 | 88 | 1 | 13 |  |
| **Breslow** | >2 mm | 12 | 50 | 10 | 83 | 2 | 17 | 0.09<sup>b</sup> |
|  | =<2 mm | 12 | 50 | 5 | 42 | 7 | 58 |  |
| **Ulceration** | Yes | 7 | 39 | 5 | 71 | 2 | 29 | 0.37<sup>b</sup> |
|  | No | 11 | 61 | 5 | 45 | 6 | 55 |  |
| **Mutational status** | BRAF | 15 | 54 | 7 | 47 | 8 | 53 | 0.33<sup>b</sup> |
|  | NRAS | 5 | 18 | 4 | 80 | 1 | 20 |  |
|  | None | 8 | 29 | 6 | 75 | 2 | 25 |  |
| **Mitotic index** | Median (IQR) | 7 (10) |  | 10 (8.5) |  | 4.5 (11.75) |  | 0.79<sup>a</sup> |
| **Checkpoint inhibitor treatment** | Yes | 22 | 76 | 13 | 59 | 9 | 41 | 1<sup>b</sup> |
|  | No | 7 | 24 | 4 | 57 | 3 | 43 |  |
| **Survival** | Alive | 13 | 45 | 5 | 38 | 8 | 62 | 0.07<sup>b</sup> |
|  | Dead | 16 | 55 | 12 | 75 | 4 | 25 |  |

<sup>a</sup>Two sample Student's t-test <sup>b</sup>Fisher's Exact

Patients were grouped based on whether they developed distant metastasis (DM; n=17) or not (noDM; n=12) within the shortest follow-up available for patients without DM and still alive at study conclusion (25.1 months; median follow-up of 33.5 months; Supp. Fig. S3). By assessing the overall % of melanoma and CD45^+^ immune cells, no differences between the DM and noDM groups were observed (Fig. 1c). A high heterogeneity was found in both groups, with tumors containing from 8.5 to 92.9 % CD45^+^ cells. Within the CD45^+^ immune cells, a trend, though not significant, towards higher proportions of CD8^+^ T cell and lower fractions of CD19^+^ B cell were observed in the noDM group compared to the DM group (Fig. 1d).

### No association between melanoma phenotype and DM

No significant differences were found in the proportion of cells expressing the established melanoma markers AXL and Melan-A (Fig. 2a, Supp. Fig. S4a) or in correlation between their expression intensities (Supp. Fig. S4b) when comparing the two groups. To take advantage of the high dimensional data, FlowSOM was used for unsupervised cluster identification across the CD45^-^CD3^-^ melanoma cells. This allowed for the identification of eight distinct cell populations within the melanoma cell cluster (Fig. 2b). Each of these subpopulations were characterized by difference in expression levels of distinct cellular markers (Fig. 2c). While large heterogeneity was observed across the samples, no significant differences were found when comparing noDM and DM groups (Fig. 2d).

**Figure 2.**
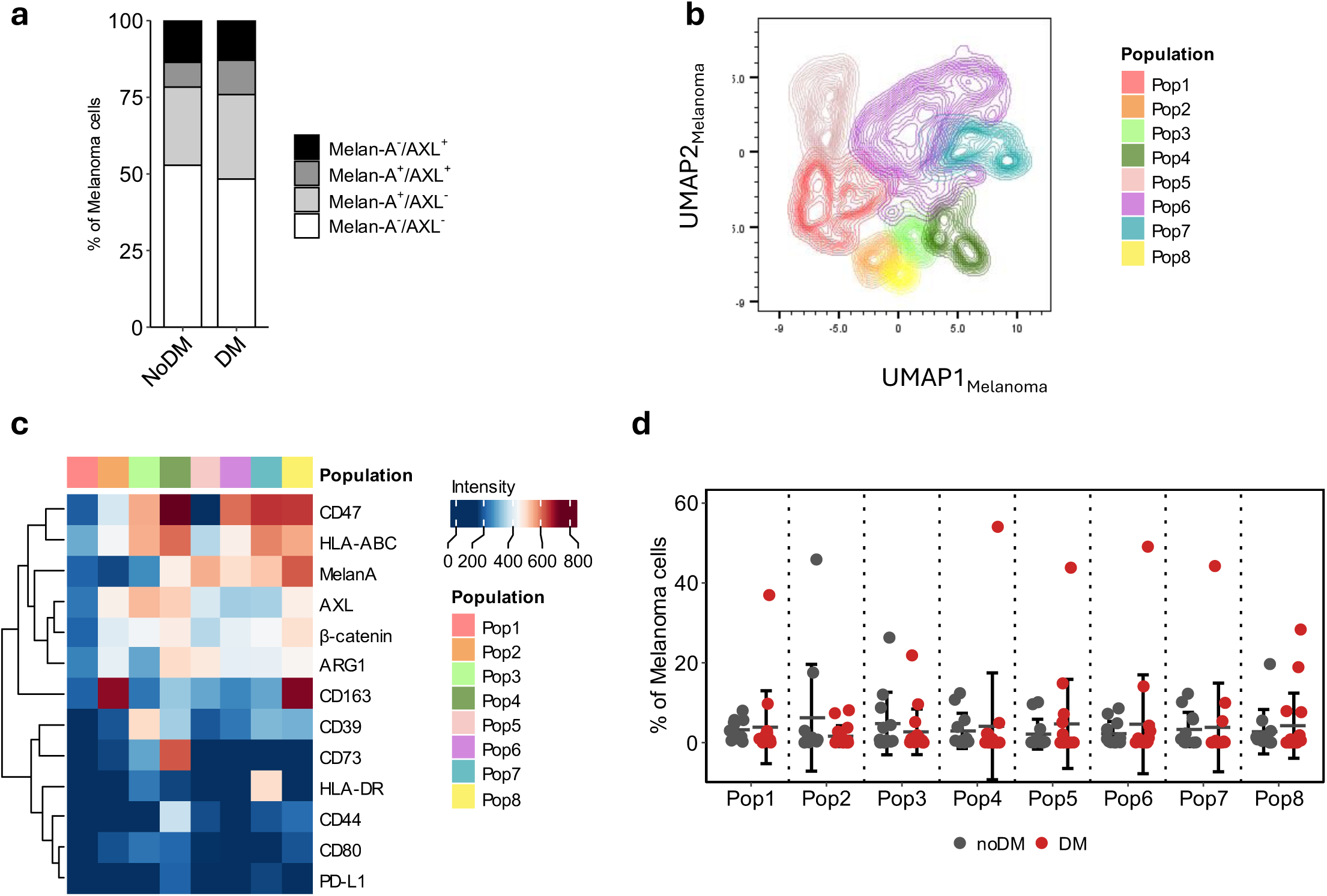
Melanoma phenotype is not associated with distant metastasis. **a.** Mean proportion of Melan-A^-^/AXL^+^, Melan-A^+^/AXL^+^, Melan-A^+^/AXL^-^, Melan-A^-^/AXL^-^ across all biopsies. **b.** UMAP of melanoma subcluster highlighting Population 1-8 identified by FlowSOM. **c.** Populations in b clustered by their expression of cell markers used for FlowSOM analyses. **d.** Populations/Clusters from b are shown as the relative proportion of melanoma cells represented by each sample.

### Trend towards higher PD-1 and activated T cell compartments in patients not developing distant metastases

Since patients in this study cohort were subsequently treated with adjuvant αPD-1, we sought to evaluate whether the expression of PD-1 in tumor biopsies was associated with progressive disease. PD-1 expression was mainly found on cells in the T cell compartment (CD8^+^ or CD4^+^; Supp. Fig. S1). We found a trend towards higher fraction of PD-1^+^ cells in the noDM group across all CD45^+^ cells, as well as CD8^+^ and CD4^+^ T cells (Fig. 3a). As expression intensity is an important measure of downstream signaling, we assessed the geometric mean intensity (GMI), and as expected, observed a similar trend towards higher PD-1 expression in the noDM group for CD45^+^ cells, as well as the distinct T cell subsets (Fig. 3b). Strong positive correlation between PD-1^+^ fractions and expression intensities was observed for CD45^+^ immune cells (Fig. S4c), CD8^+^ T cells (Fig. 3c) and CD4^+^ T cells (Fig. S4d).

**Figure 3.**
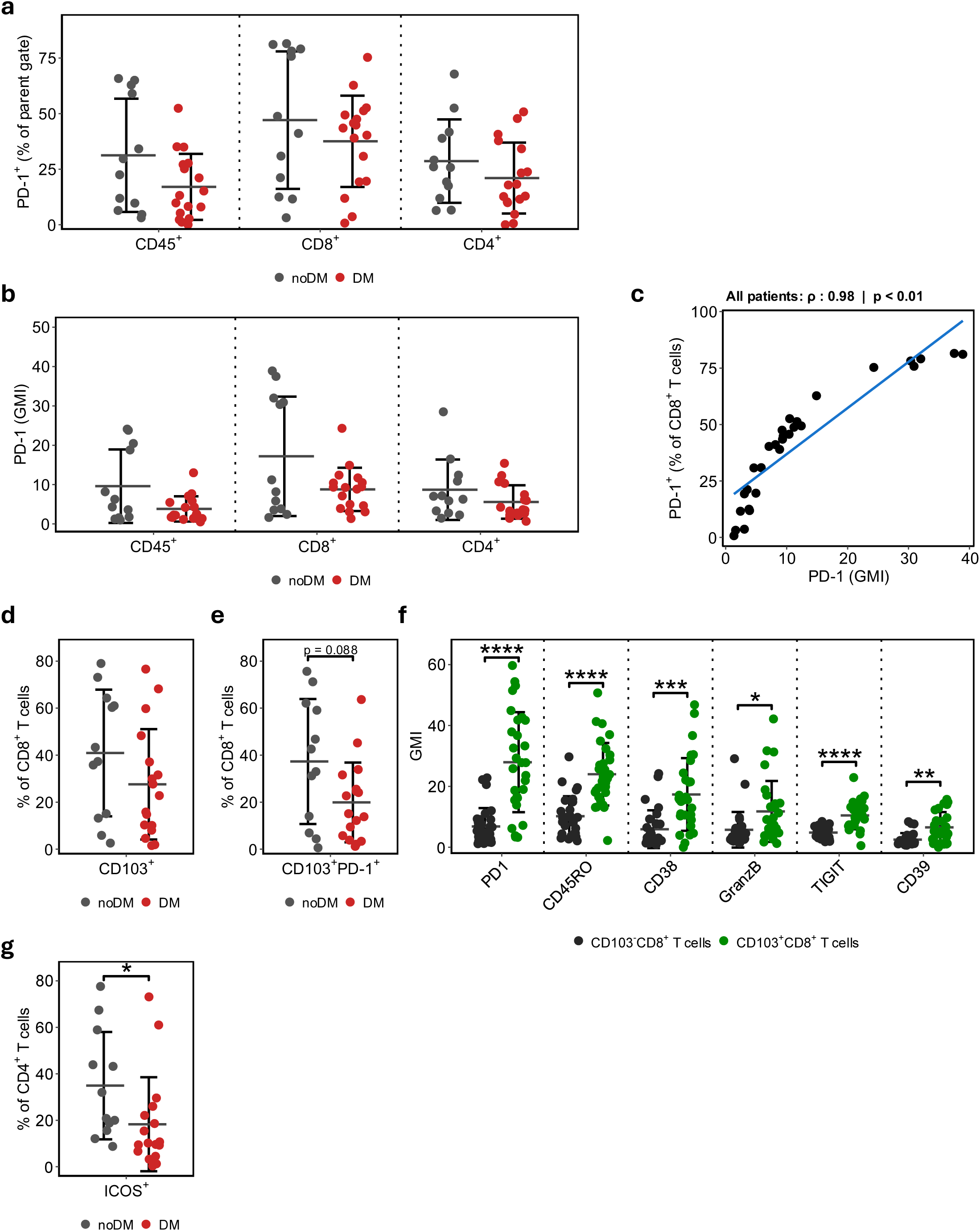
Biopsies from noDM patients are increased in activated phenotypes of CD8^+^ and CD4^+^ T cells. **a.** Fraction of PD-1 positive cells within the CD8^+^ and CD4^+^ T cell populations, as well as across all CD45^+^ cells. PD-1 is represented as the proportion of positive cells in the respective population. **b.** GMI for PD-1 in the same population as a). **c.** Spearman correlation of PD-1^+^ proportion and intensity (GMI) in CD8^+^ T cells across all samples. **d.** Proportion of CD103^+^ within the CD8^+^ T cell population. **e.** GMI of surface markers in the CD103^+^ and CD103^-^ CD8^+^ T cell populations. **f.** Proportion of CD103^+^PD-1^+^ cells within the CD8+ T cell population. **g.** Proportion of ICOS^+^ cells within the CD4+ T cell population. Proportion of positive cells or GMI shown as mean±SD.

We further assessed whether the slightly higher fraction of CD8^+^ T cells in the noDM group (Fig. 1d) was associated with changes in immune cell activation status and checkpoint expression beyond PD-1. Tissue-resident memory (T_RM_) cells have been associated with improved survival in immunotherapy-naïve patients and expand significantly during anti-PD- 1 treatment (15). In line with this observation, a trend towards a higher fraction of T_RM_ defined by CD103^+^CD8^+^ expression was found in the noDM samples (Fig. 3d; p = 0.166). Interestingly, the proportion of PD-1 expressing T_RM_ was close to two times higher in the noDM group (Fig. 3e; p = 0.088). Additionally, the T_RM_ population demonstrated significant increase in expression of markers CD45RO, CD38, Granzyme B, TIGIT and CD39 (Fig. 3f and Supp. Fig. S4f), as well as minor increase in HLA-DR, TIM-3 and LAG-3 (Supp. Fig. S4e-f), compared to the CD103^-^CD8^+^ T cells, supporting an activated immune phenotype. Similarly, a significantly higher fraction of activating marker ICOS^+^ (p = 0.018) was observed on CD4^+^ T cells in the noDM group (Fig. 3g). Altogether, these data suggest enrichment of an anti-tumoral microenvironment with activated T_RM_ and CD4^+^ T cells in the RLN from patients with noDM compared to DM.

### CD38^hi^CD19^dim^ cells are associated with improved distant metastasis-free survival

No significant differences were found in CD19^+^ B cells between the DM and noDM groups, despite a trend towards fewer CD19^+^ B cells in noDM samples (Fig. 1d). Interestingly, by examining the “Other immune” cluster, a small fraction of CD38^+^ cells with intermediate levels of CD19-expression (labeled CD38^hi^CD19^dim^) were identified (Fig. 4a, Supp. Fig. 5a).

**Figure 4.**
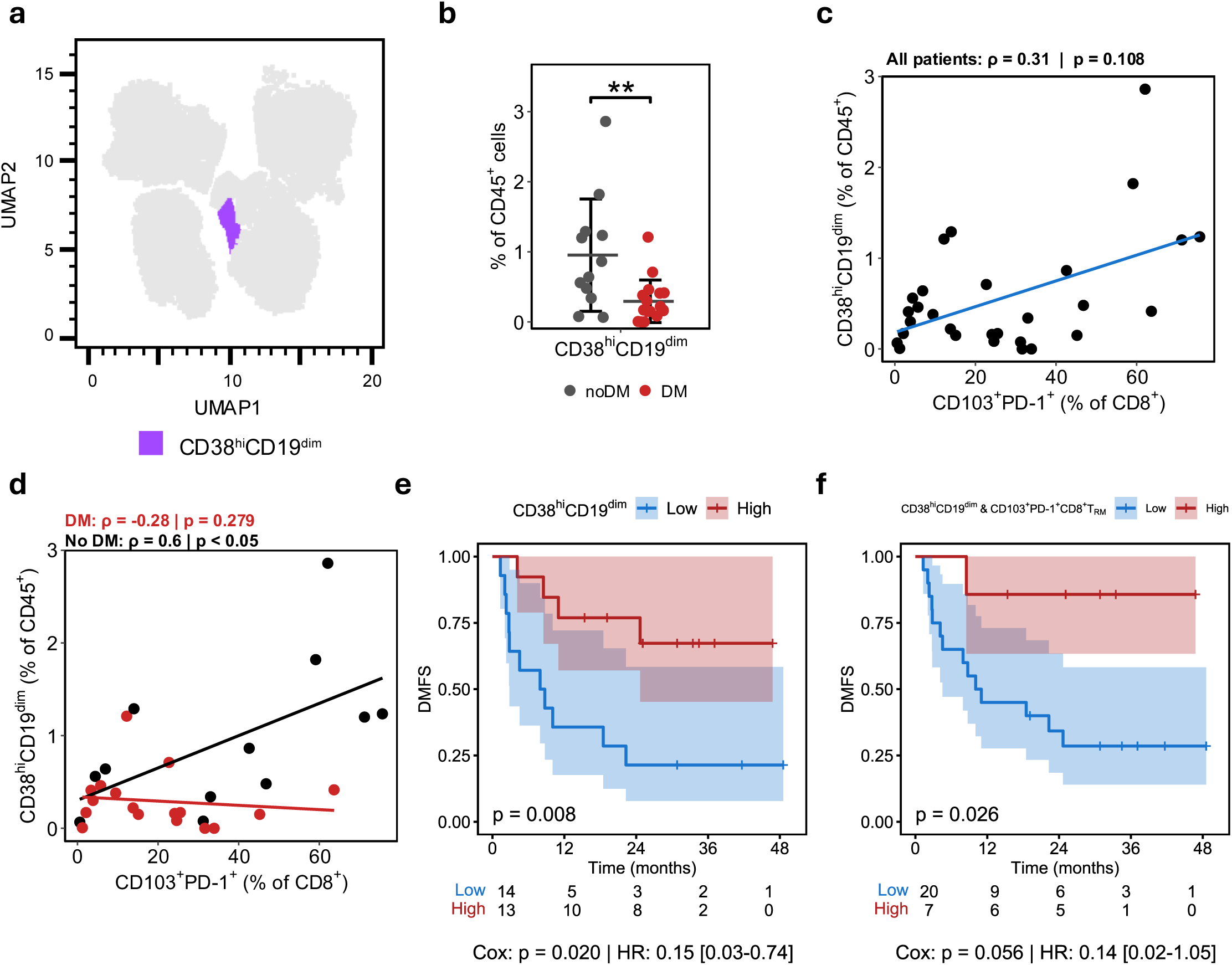
Reduced fraction of CD38^hi^CD19^dim^ associated with distant metastasis. **a.** The cell population of CD19^dim^ cells with a corresponding increase in CD38 are highlighted in purple. **b.** Proportion of CD38^hi^CD19^dim^ in DM and noDM groups relative to CD45^+^ cells. Mean±SD. **c-d.** Spearman correlation of fraction of CD38^hi^CD19^dim^ and CD103^+^PD-1^+^CD8^+^ T cells, across all samples (**c**) and separated on DM-status (**d**). **e.** Kaplan-Meier plot showing distant metastasis-free survival (DMFS) in patients stratified on fraction of CD38^hi^CD19^dim^ higher (red) or lower (blue) than the median. **f.** Kaplan-Meier plot stratified on fraction >median in both CD38^hi^CD19^dim^ and CD103^+^PD-1^+^CD8^+^ T_RM_ cells, as *High*, remaining samples *Low*. P- values in e and f are calculated by log-rank test, while p-value for Cox-regression and corresponding Hazard ratio [95 % CI] are displayed at the bottom of each plot.

By comparing the DM-groups, a significantly higher fraction of CD38^hi^CD19^dim^ was found in patients without DM (Fig. 4b). Only moderate correlation between CD38^hi^CD19^dim^ and the CD103^+^PD-1^+^CD8^+^ T cells were found (Fig. 4c). However, when separating samples by DM-status a strong positive correlation was found between the cell types only in the noDM patients (Fig. 4d). Similarly, inverse correlation between CD38^hi^CD19^dim^ and CD19^+^ B cells were found only in tumors from the noDM group (Supp. Fig. S4b-c), while no strong correlation was found between CD38^hi^CD19^dim^ and ICOS^+^CD4^+^ T cells in either of the DM groups (Supp. Fig. S5d-e).

To further assess whether these differences could be linked to lack of progression and potential long-term benefit of ICI in these patients, we performed survival analyses defining CD38^hi^CD19^dim^ *High* and *Low* groups by dividing at the median expression (Fig. 4e). The *High* expression group showed significantly improved distant metastasis-free survival (DMFS) compared to the *Low* group (log-rank p-value = 0.008), with an area under the curve (AUC) of 0.81 and specificity >90 % (Supp. Fig. 5f). Additionally, Cox-regression was significant (p-value = 0.02) with a hazard ratio (HR) of 0.15 (95 % CI: 0.03 – 0.74), indicating more than 5-fold reduction in risk of distant metastasis for patients in the *High* group. Stratifying patients on the CD103^+^PD-1^+^CD8^+^ subpopulation, did only define a *High* expressing group doing marginally, though not significantly, better (Supp. Fig. S5g; p=0.093). However, when making a surrogate for both markers (*High* group defined as >median expression of both CD38^hi^CD19^dim^ and CD103^+^PD-1^+^CD8^+^ T cells), we identified a small group having excellent outcome with only one distant metastasis during 4-year follow up (p-value = 0.026; Fig. 4f).

### Loss of B cell score and CD38 expression associated with poor prognosis in stage III melanoma patients receiving no adjuvant therapy

To further evaluate the prognostic impact of CD38^hi^CD19^dim^ and CD103^+^PD-1^+^CD8^+^ T cells, we utilized a NanoString dataset generated in-house from 125 regional lymph node biopsies collected from patients with stage III melanoma treated with surgery only. We first assessed how the tumor-infiltrating lymphocyte (TIL) score generated by pathological evaluation was associated with survival in this patient cohort. Interestingly, patients with Hot tumors (TIL score > median) did not show improved survival compared to patients with Cold tumors (Fig. 5a). By using the cell subtype specific gene signatures provided with the IO360 panel, we proceeded to infer both CD8^+^ T cell and B cell scores, representing the two major lymphocyte populations found in the CyTOF cohort. No correlation was found between the CD8^+^ T cell and TIL scores (Fig. 5b), while strong correlation was found between B cell and TIL scores (Fig. 5c). By stratifying using the CD8^+^ T cell score inferred by NanoString we found improved survival in patients with high CD8^+^ T cell score (p = 0.033; Fig. 5d), but not when stratified by B cell score (Supp. Fig. S6a). By revisiting the cut-off for TIL infiltration, we found a subset of patients with <2.5 % TILs with reduced time to distant metastasis (p = 0.0018; Fig. 5e). This “*Ultra-cold*” group was associated with a highly significant drop in B cell score, but not in CD8^+^ T cell score (Fig. 5f). As the B cell score in a bulk setting will represent most B cell subsets, including plasmablast-like cells, we further analyzed the expression of selected markers linked to plasmablast-like and CD8^+^ T cell activation in this group. While the *Ultra-cold* group had significantly reduced expression of markers associated with plasmablast-like cells; *CD38*, *CD27*, *CD19* and *CD79A/B*, no significant changes were seen for the genes *PDCD1* and *ITGAE,* encoding PD-1 and CD103 respectively (Supp. Fig. S6b). Notably, *CD38* and *CD19* expression were strongly correlated only in patients with TIL <2.5 % (Supp. Fig. S6c), compared to *CD38* and *CD8A* that showed similar correlation in the two groups (Supp. Fig. S6d).

**Figure 5.**
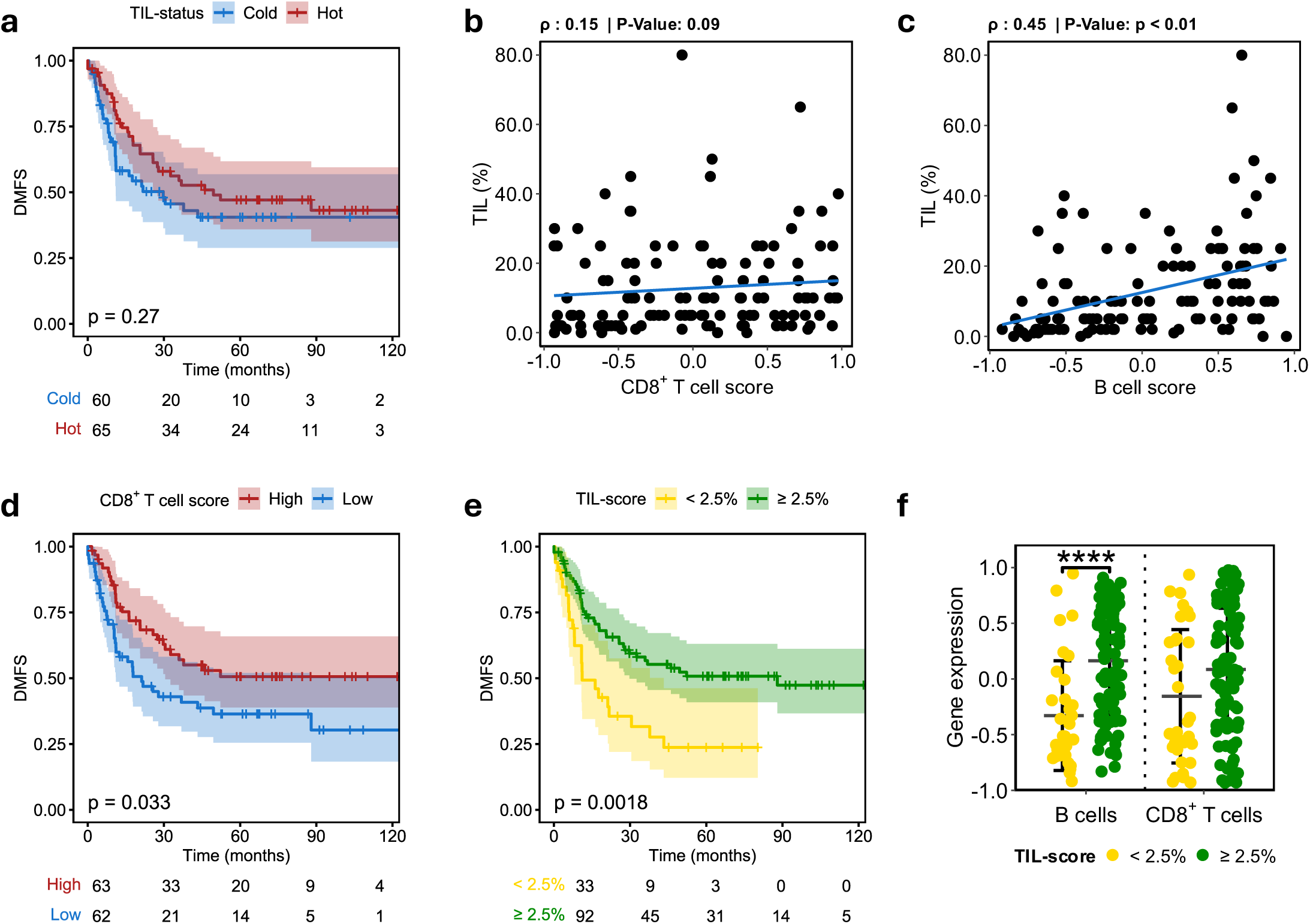
Low tumor-infiltrating lymphocytes associated with reduced B cell score and poor outcome. **a.** Kaplan-Meier plot showing distant metastasis-free survival (DMFS) in patients stratified on tumor-infiltrating lymphocytes (TIL), separated into Cold (<median infiltration) and Hot (>median infiltration). **b-c.** Spearman correlation of fraction of TIL and CD8^+^ T cell (**b**) and B cell (**c**) gene scores from NanoString. **d.** Kaplan-Meier plot showing DMFS in patients stratified on CD8^+^ T cell gene score from NanoString, separated by the median into *Low* and *High*. **e.** Kaplan-Meier plot showing DMFS in patients stratified by ≥ 2.5 % pathologist TIL score. **f.** B cell and CD8^+^ T cell gene score from NanoString in groups stratified by ≥ 2.5 % pathologist TIL score. Mean±SD. P-values in a, d and e were calculated by log-rank test.

## Discussion

In this study we have identified a subpopulation of CD38^hi^CD19^dim^ cells present at baseline, that were associated with improved distant metastasis-free survival in melanoma patients receiving adjuvant treatment with αPD-1. Additionally, the higher fraction of CD38^hi^CD19^dim^cells was strongly correlated with increased proportion of CD103^+^PD-1^+^CD8^+^ T_RM_ cells in patients with noDM. We hypothesize that these subpopulations together generate an immune microenvironment beneficial for effect of αPD-1 treatment. Evaluation of these subpopulations as predictive biomarkers of response to αPD-1 blockade could aid in selection of patients with benefit of ICI.

Even though melanoma cells were frequent in our cohort, we did not find specific phenotypes associated with disease progression and reduced response to ICI (Fig. 1a and 2). The relative small cohort could partly explain this observation, however, melanoma is known to be heterogenous, suggesting that the spatial context of the tumor clones could be necessary to fully understand the phenotypes associated with treatment response or absence thereof (16). Similarly, only minor trends were seen in the major immune cell populations, as CD4^+^ and CD8^+^ T cells showed a slight decrease and CD19^+^ B cells a slight increase in patients that developed DM (Fig. 1d). This is in line with previous studies in metastatic melanoma, where the presence of CD8^+^ T cells has been reported in lesions responding to ICI (17).

In cutaneous melanoma, stem-like phenotypes are associated with metastatic spread and treatment resistance (18). This phenotype is often represented by high expression of the receptor tyrosine kinase AXL and low expression of melanocytic markers like Melan-A (19, 20). In line with the observation in the present cohort (Fig. 2a, Supp. Fig. S4a-b), we have previously shown how AXL is heterogeneously expressed in lymph node metastases from stage III melanoma (21), as wells as in stage IV disease (22). Melanoma cells were mainly defined by CD45^-^CD3^-^, thus other cell types could potentially be assigned to this cluster.

However, unsupervised analysis with FlowSOM did not identify subclusters associated with DM based on the markers available in our CyTOF panel. Altogether, this suggests that larger cohorts, in addition to spatial assessment, of RLN biopsies would be necessary for further identification of melanoma phenotypes associated with development of distant metastasis. Such studies would allow investigation of melanoma cell plasticity in a spatial context and may unveil a spectrum of activation states central in treatment escape and identify microenvironmental factors critical for response to ICI.

While ICI has been highly successful in melanoma (3), there is still a lack of reliable biomarkers for predicting benefit from therapy. Recent efforts utilizing -omics approaches have shown some promise for primary melanoma (11), while predictive biomarkers for stage III disease are still lacking. In a recent study, genes previously associated with ICI response (12) were used to generate an IFNg-based signature with prognostic value in stage III melanoma, but with limited predictive value of response to ICI (10). PD-L1 expression, evaluated through immunohistochemistry, is routinely used as biomarker for selecting PD- 1/PD-L1 inhibitors in various cancers. However, in melanoma, this practice is less common due to considerable response rates even in patients with PD-L1 negative tumors. Notably, the expression of PD-L1 has been shown to increase in melanoma LN metastasis compared to primary tumors (23). In our study, no significant association was found between PD-1 expression on CD45^+^ immune cells and development of distant metastasis in patients receiving αPD-1 (Fig. 3a-c).

Several studies have highlighted the importance of different activation states of CD8^+^ T cells as predictors of response to immune therapy (6, 15, 24, 25). In the present study we identified a higher proportion of CD103^+^CD8^+^ T cells, generally referred to as CD8^+^ T_RM_ cells, characterized by high expression of the CD103 surface marker (26). CD8^+^ T_RM_ has previously been found at increased frequency in melanoma lymph node metastasis that do not develop into distant metastasis (6, 15). We found that CD8^+^ T_RM_ activation is characterized by increased expression of several activation markers (Fig. 3g), including CD39, which has been associated with a favorable response to ICI when upregulated on CD8^+^ T_RM_ (25). Additionally, targeting the increased expression of surface markers such as PD-1, CD38 and TIM-3 found on this population has been suggested for improving ICI sensitivity (15, 27–29). Altogether, this illustrates that high pre-treatment proportion of activated T_RM_ with cytotoxic features generates a favorable microenvironment for effect of ICI therapy.

Interestingly, we found that CD8^+^ T cell gene score, but not pathologist-generated TIL score, is associated with good prognosis in our no-adjuvant cohort (Fig. 5). TIL score itself seemed to most accurately represent B cell infiltration rather than CD8^+^ T cells. Lower B cell infiltration was also seen in the CyTOF data (Fig. 1d) in patients with noDM after ICI, but B cells score was not directly linked to prognosis in the NanoString cohort. Others have observed CD19^+^ B cells to be associated with good outcome in stage III melanoma, but it is worth noting that their study was performed in stage III melanoma collected from skin and RLN (30) representing a slightly different cohort than we have used in this study. Reports indicate impaired B cell activation to serve as a step in development of advanced melanoma (31), and clonal B cell receptor (BCR) repertoire to be good indicators of response to ICI in metastatic melanoma (32). Altogether this suggests TIL score on its own to be of limited use when assessing the tumors immune-status in the context of likely ICI response, as both T cell and B cell activation states are important factors for accurately assessing the immune-activation.

We identified a small population of CD38^hi^CD19^dim^ cells strongly associated with good prognosis after ICI treatment. This subpopulation showed marker expression in line with those of plasmablast-like cells (33), having high expression of CD38, low expression of CD19, and absence of T cell lineage markers (Supp. Fig. S5a). Interestingly, increased proportion of these plasmablast-like cells before ICI strongly correlated with longer time until distant metastasis and showed improved prognostic impact compared to the PD-1^+^ CD8^+^ T_RM_ fraction alone (Fig. 4e, Supp fig. S5g). Patients with high fractions of both PD-1^+^ CD8^+^ T_RM_ and CD38^hi^CD19^dim^ showed an even better outcome with only 1/7 patients developing distant metastasis. In addition, the fraction of these subtypes showed strong correlation in samples from patients that did not develop DM (Fig. 4d), but not in patients that did develop DM, indicating a potential importance of both these cell types in defining a favorable microenvironment for response to ICI. In line with our observation, Griss *et al*. observed plasmablast-like (CD38^+^CD19^+^) cells to be central in shaping the microenvironment in cutaneous melanoma, and to be associated with increased PD-1^+^ T cell activation after anti-PD-1 blockade *in vitro* (33). Additionally, they found that higher frequency of these plasmablast-like cells predicted response to immune checkpoint blockade and survival in metastatic melanoma (stage IV), supporting the presence of an anti-tumoral immune microenvironment reinvigorated by ICI therapy. The tumor-induced plasmablast-like-enriched B cell population (TIPB) signature defined in the Griss study, consisting of *CD27*, *CD38*, and *PAX5*, needs to be refined in a bulk tumor setting. For instance, while CD38 is a prominent marker on plasmablast-like cells, it is also expressed in multiple cell types, which can influence its overall expression in bulk samples. Notably, high proportion of dysfunctional PD-1^+^CD38^hi^ CD8^+^ T cells have been reported as both a predictive and therapeutic biomarker for anti-PD-1 treatment (29). This underscores the importance of accounting for the phenotype and activation status of various immune cells when defining novel biomarkers, and highlights the advantage of using single cells rather than bulk signatures as prognostic markers. Furthermore, Griss *et al* reports strong correlation between the TIPB and multiple signatures of T cells and cytotoxic activity, suggesting that the TIPB is significantly influenced by CD8^+^ T cell expression. Using NanoString gene expression data from a cohort of untreated patients, we aimed to assess expressional changes in genes associated with plasmablast-like genes and B cell scores. While, in line with the CyTOF data, high CD8^+^ T cells score were associated with improved survival in patients not receiving adjuvant therapy, B cells on its own was not associated with disease prognosis. Interestingly, a subset of *Ultra-cold* tumors with reduced lymphocyte infiltration (TIL < 2.5 %) was strongly correlated with worse outcome and significant reduction in B cell scores. This was accompanied by reduced expression of both *CD38* and *CD27*, genes central for the plasmablast-like phenotype, but not reduction in CD8^+^ T cell score. We hypothesize that the absence of plasmablast-like cells is central in the poor prognosis of this group. Along this line, we found strong correlation between *CD19* and *CD38* only in this subset of patients, suggesting the drop in *CD38* to come from changes in B cell presence rather than CD8^+^ T cells, highlighting the potential for using plasmablast-like cells as stratification criteria in stage III melanoma. Our study demonstrates how single cell approaches could enhance stratification of these patients relative to bulk approaches.

While this study identifies several cell types associated with favorable prognosis in patients receiving adjuvant αPD-1 therapy, it is important to highlight that without any treatment response measure, we are unable to conclude whether these features can directly predict response to αPD-1 treatment. Notably, we observed similar associations between CD8^+^ T cell scores, genes associated with plasmablast-like cells and good prognosis among patients not receiving adjuvant treatment in the NanoString cohort. This could suggest that enrichment of these cells prior to therapy in general is indicative of a better outcome, regardless of adjuvant treatment. Additionally, the current standard-of-care for stage III melanoma includes neo- adjuvant αPD-1 therapy prior to lymph node resection. Collecting appropriate specimens in this context could help evaluate whether T_RM_ and plasmablast-like cells serve as predictive markers for αPD-1 treatment according to the current standard-of-care, while also identifying patients with favorable prognoses, as the current study suggests.

## Conclusion

In this study, several immune cell types in the RLN were found associated with better prognosis in stage III melanoma treated with adjuvant αPD-1 after lymph node surgery. Notably, high fractions of CD38^hi^CD19^dim^ plasmablast-like cells and activated CD103^+^PD- 1^+^CD8^+^ T_RM_ were strongly associated with distant metastasis-free survival. While the findings in this explorative study require validation in an independent patient cohort, it highlights the potential for using immune cell subsets as prognostic biomarkers, and possibly for predicting response to αPD-1 in this patient group.

## Additional Information

## Acknowledgements

We would like to thank patients with metastatic melanoma treated at the Norwegian Radium Hospital between 2020 - 2022 for their participation in this research study. Huge thanks to Idun Dale Rein, Monica Bostad and Heidi Ødegaard Notø at the Flow cytometry and Pre-Clinical Imaging Core Facility for excellent technical support with CYTOF instrument and analysis (Institute for Cancer Research, Oslo University Hospital). We would also like to thank Martina Skrede, Marina Juraleviciute, Renate Slind Olsen and Ingrid Øverlie Bakka for technical assistance.

## Author contributions

IA, HJ, JAK, GMM and EVE drafted the manuscript. TR, RH and LF are plastic surgery specialists and secured material for research, while AVB did the pathological evaluation, and IA, MF, MN, and AH were responsible for collecting updated clinical information. IA was responsible for establishing the CyTOF protocol and performing the experiments, while IA, SKC, AP, and EVE performed downstream CyTOF analysis. BJ and EVE performed all transcriptional analysis. LP and EMI participated in data interpretation. IA, VAF, JAK, GMM and EVE conceptualized, designed, and coordinated the study, secured funding, and were involved in interpretation of the data. All authors contributed to revision, editing, and finalizing the manuscript.

## Ethics approval

Written informed consents were retrieved from all patients, and the study was approved by the Regional Ethical Committee of South-East Norway (REK#2012/2309 and REK# 2015/2434).

## Data availability

All clinical material and data presented herein are available from the authors upon reasonable request.

## List of Abbreviations

ICI: Immune checkpoint inhibitors
αPD-1: anti-PD-1 treatment
RLN: Regional lymph node
FBS: Fetal bovine serum
CyTOF®: Cytometry by Time-Of-Flight
FlowSOM: Flow Self-Organizing Map
FFPE: Formalin-fixed paraffin-embedded
IO360: nCounter PanCancer
IO 360™: Panel DM Distant metastasis
noDM: No distant metastasis
T_RM_: Tissue-resident memory CD8^+^ T cells
DMFS: Distant metastasis-free survival
AUC: Area under the curve
HR: Hazard ratio
TIL: Tumor-infiltrating lymphocyte
BCR: B cell receptor
TIPB: Tumor-induced plasmablast-like-enriched B cell population

