## Supplemental Table and Figures for "CD38^hi^CD19^dim^ cells in lymph nodes predict favorable prognosis in patients with stage III melanoma receiving adjuvant PD-1-blockade"

### Supplementary Table S1

List of antibodies

| Target | Mass | Metal | Clone | Manufacturer | Catalog # | Final concentration | Additional Info |
|---|---|---|---|---|---|---|---|
| **CD45** | 89 | Y | HI30 | Standard BioTools | 3089003C | 1:400 | |
| **CD19** | 142 | Nd | HIB19 | Standard BioTools | 3142001C | 1:400 | |
| **ICOS** | 143 | Nd | C398.4A | Standard BioTools | 3143025C | 1:400 | |
| **HLA-ABC** | 144 | Nd | W6-32 | Standard BioTools | 3144017C | 1:400 | |
| **CD163** | 145 | Nd | GHI/61 | Standard BioTools | 3145010B | 1:200 | Intracellular |
| **CD8a** | 146 | Nd | RPA-T8 | Standard BioTools | 3146001C | 1:400 | |
| **β-catenin** | 147 | Sm | D10A8 | Standard BioTools | 3147005C | 1:400 | Intracellular |
| **CD16** | 148 | Nd | 3G8 | Standard BioTools | 3148004B | 1:200 | |
| **CD25** | 149 | Sm | 2A3 | Standard BioTools | 3149010C | 1:400 | |
| **LAG3** | 150 | Nd | 11C3C65 | Standard BioTools | 3150030B | 1:200 | |
| **CD103** | 151 | Eu | Ber-ACT8 | Standard BioTools | 3151011B | 1:200 | |
| **TCRgd** | 152 | Sm | 11F2 | Standard BioTools | 3152008B | 1:200 | |
| **TIM3** | 153 | Eu | F38-2E2 | Standard BioTools | 3153008B | 1:200 | |
| **TIGIT** | 154 | Sm | MBSA43 | Standard BioTools | 3154016B | 1:200 | |
| **PD-1** | 155 | Gd | EH12.2H7 | Standard BioTools | 3155009B | 1:200 | |
| **CD14** | 156 | Gd | HCD14 | Standard BioTools | 3156019B | 1:200 | |
| **Melan-A** | 158 | Gd | #872719 | R&D systems | MAB8008 | 1:200 | Intracellular/Conjugate |
| **CD39** | 160 | Gd | A1 | Standard BioTools | 3160004C | 1:400 | |
| **AXL** | 161 | Dy | Polyclonal | R&D systems | AF154 | 1:400 | Intracellular/Conjugate |
| **CD80** | 162 | Dy | 2D10.4 | Standard BioTools | 3162010B | 1:200 | |
| **CD33** | 163 | Dy | WM53 | Standard BioTools | 3163023C | 1:400 | |
| **Arginase-1** | 164 | Dy | 14D2C43 | Standard BioTools | 3164030B | 1:200 | Intracellular |
| **CD45RO** | 165 | Ho | UCHL1 | Standard BioTools | 3165011C | 1:400 | |
| **CD44** | 166 | Er | BJ18 | Standard BioTools | 3166001C | 1:200 | |
| **CD11b** | 167 | Er | ICRF44 | Standard BioTools | 3167011C | 1:400 | |
| **CD73** | 168 | Er | AD2 | Standard BioTools | 3168015B | 1:200 | |
| **CD45RA** | 169 | Tm | HI100 | Standard BioTools | 3169008B | 1:400 | |
| **CD3** | 170 | Er | UCHT1 | Standard BioTools | 3170001C | 1:400 | |
| **Granzyme B** | 171 | Yb | GB11 | Standard BioTools | 3171002C | 1:200 | Intracellular |
| **CD38** | 172 | Yb | HIT2 | Standard BioTools | 3172007C | 1:200 | |
| **HLA-DR** | 173 | Yb | L243 | Standard BioTools | 3173005C | 1:400 | |
| **CD4** | 174 | Yb | SK3 | Standard BioTools | 3174004C | 1:400 | |
| **PD-L1** | 175 | Lu | 29E.2A3 | Standard BioTools | 3175017B | 1:200 | |
| **CD56** | 176 | Yb | N901 | Standard BioTools | 3176009C | 1:200 | |
| **CD47** | 209 | Bi | CC2C6 | Standard BioTools | 3209004C | 1:200 | |

**Supplementary Fig. S1.** Marker expression

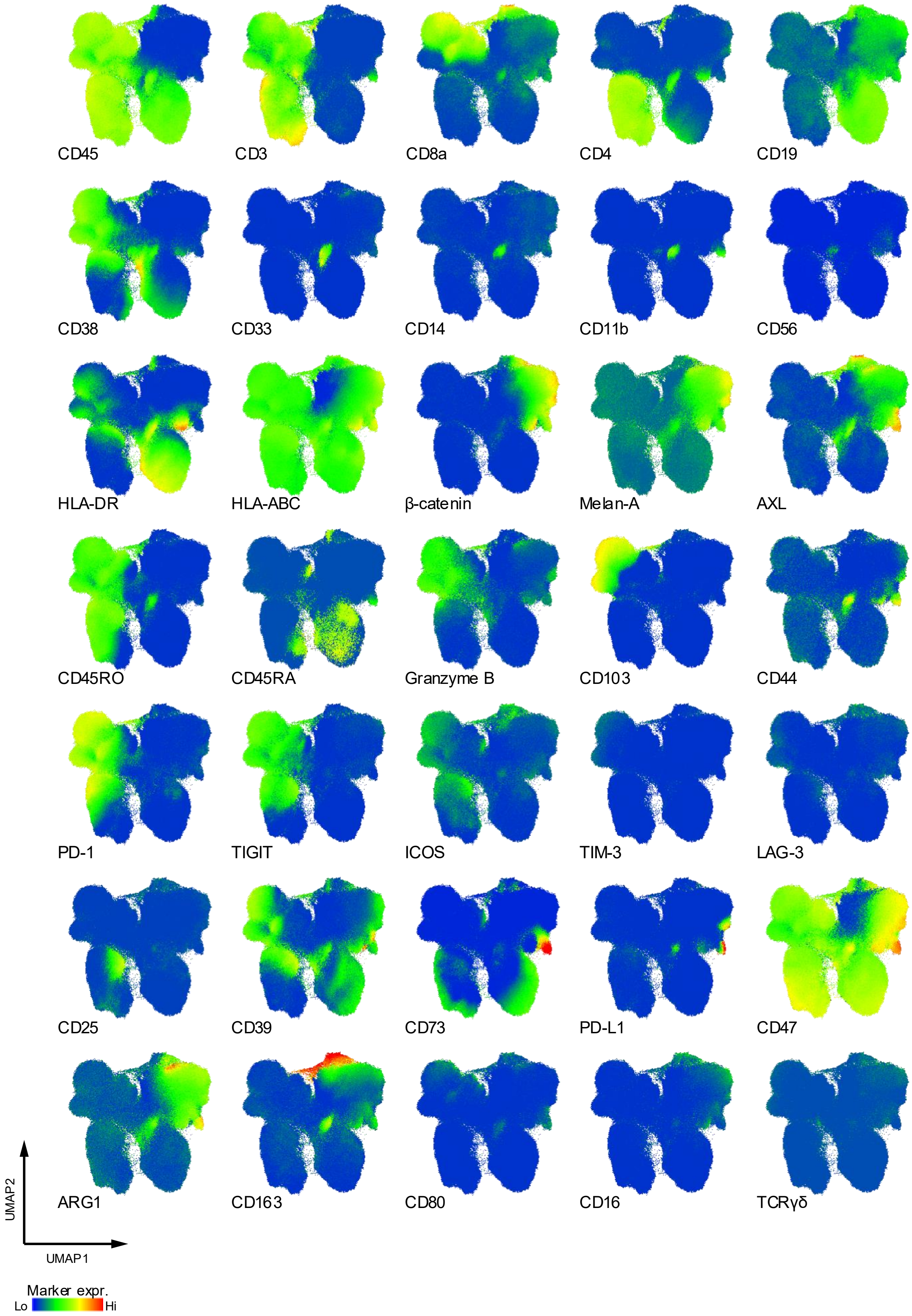


**Supp. Fig. S1.** *Marker expression.* UMAP indicating the expression levels of all markers included in the CyTOF panel.

**Supplementary Fig. S2.** Gating chart

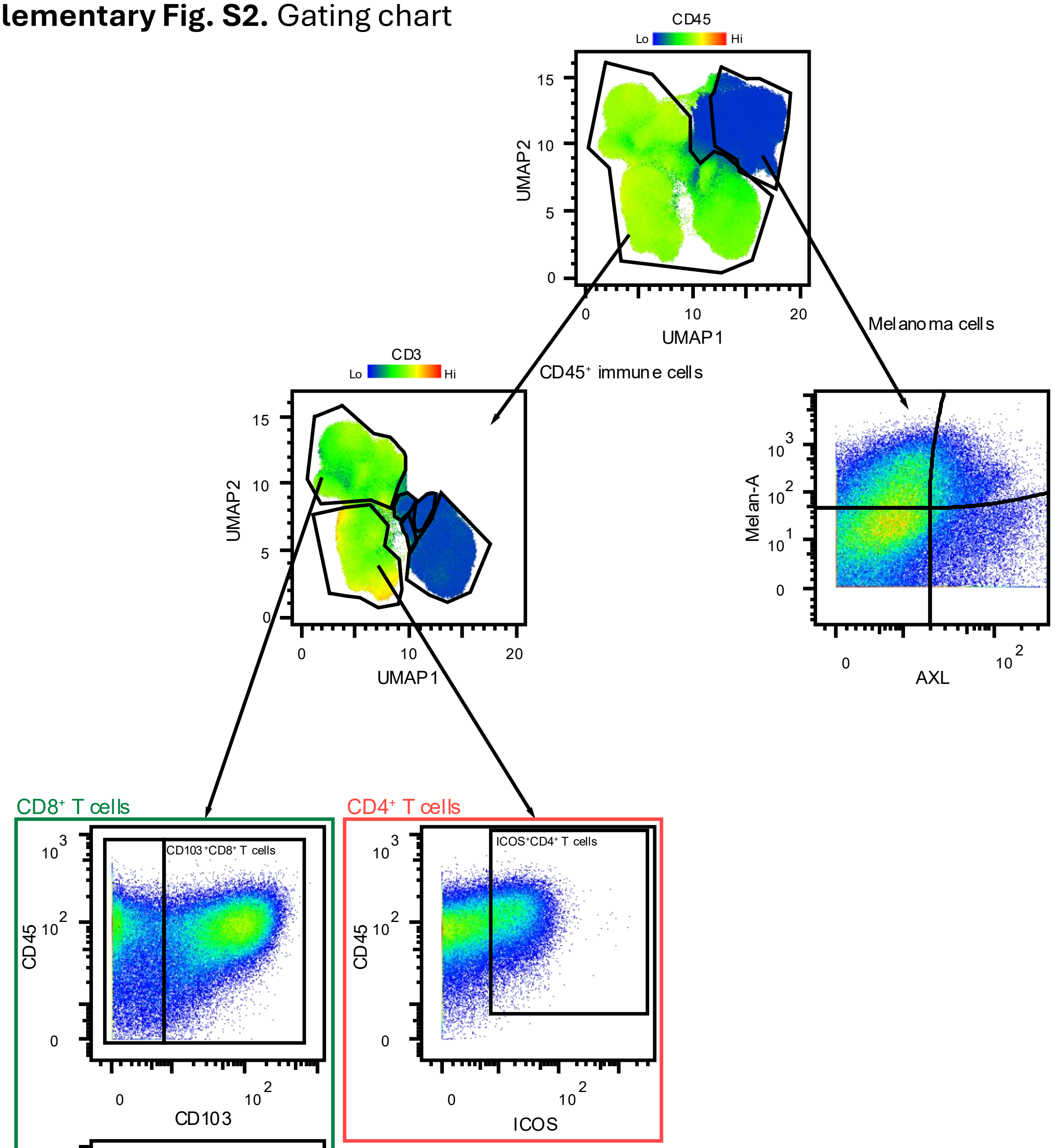


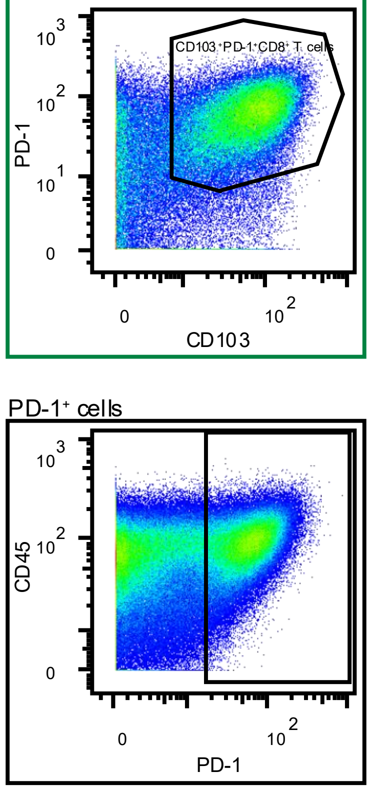


Melanoma: $CD45^{-}CD3^{-}AXL^{+/-}Melan\text{-}A^{+/-}\beta\text{-}catenin^{+/-}$

$CD8^{+}$ T cells: $CD45^{+}CD3^{+}CD19^{-}CD8^{+}$

PD-$1^{+}CD8^{+}$ $T_{RM}$: $CD45^{+}CD3^{+}CD19^{-}CD8^{+}CD103^{+}PD\text{-}1^{+}$

$CD4^{+}$ T cells: $CD45^{+}CD3^{+}CD19^{-}CD4^{+}$

Myeloid cells: $CD45^{+}CD3^{-}CD19^{-}CD33^{+}$

$CD19^{+}$ B cells: $CD45^{+}CD3^{-}CD19^{+}$

Plasmablast-like cells: $CD45^{+}CD3^{-}CD73^{-}CD38^{hi}CD19^{dim}$

**Supp. Fig. S2.** *Gating chart*. Gating of main cell population was done based on UMAP and expression on linage markers as shown in Supp Fig. S1. Subsets within the main clusters were manually gated as illustrated, with gating used for all PD-$1^{+}$ subsets showed at the bottom.

**Supplementary Fig. S3.** *Follow-up time.*

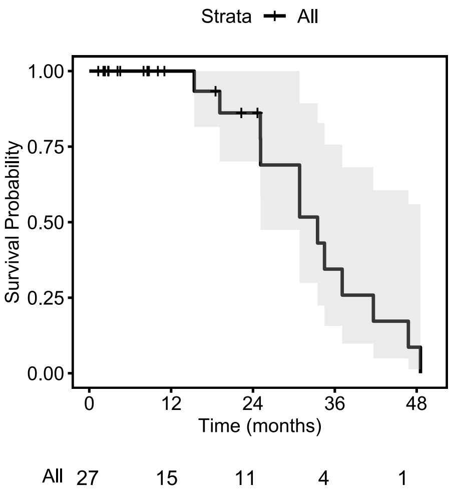


**Supp. Fig. S3.** *Follow-up time.* Reverse Kaplan-Meier survival curve showing loss to follow-up. Event represents loss to follow-up, while censoring represents distant metastasis.

**Supplementary Fig. S4.** *Melanoma and immune subpopulations.*

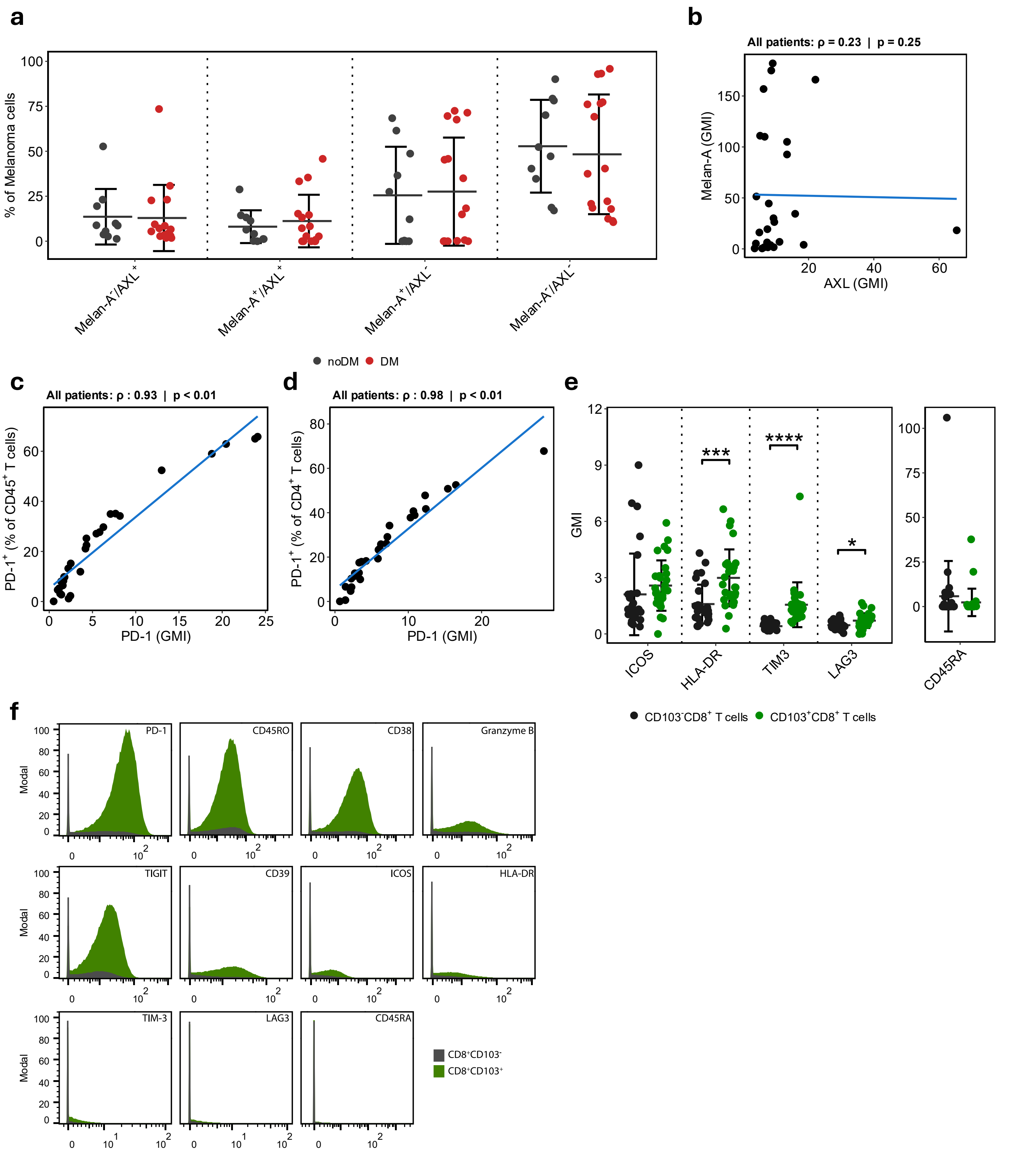


**Supp. Fig. S4.** *Melanoma and immune subpopulations in RLN biopsies.* **a.** Relative proportion of Melan-A and AXL positivity in melanoma cells as related to Figure 2a. **b.** Correlation between expression intensity as geometric mean intensity (GMI) of Melan-A and AXL in melanoma cells. **c.** Correlation between PD-1$^{+}$ fractions and GMI in CD4$^{+}$ T-cells (**c**), and CD45$^{+}$ (**d**) cell populations. **e.** Expression of low intensity markers in CD103$^{+}$ and CD103$^{-}$ CD8$^{+}$ T cells, represented as mean GMI±SD. **f.** Representative plots of relative marker expression across the CD103$^{+}$ (green) and CD103$^{-}$ (black) CD8$^{+}$ T cells.

### Supplementary Fig. S5

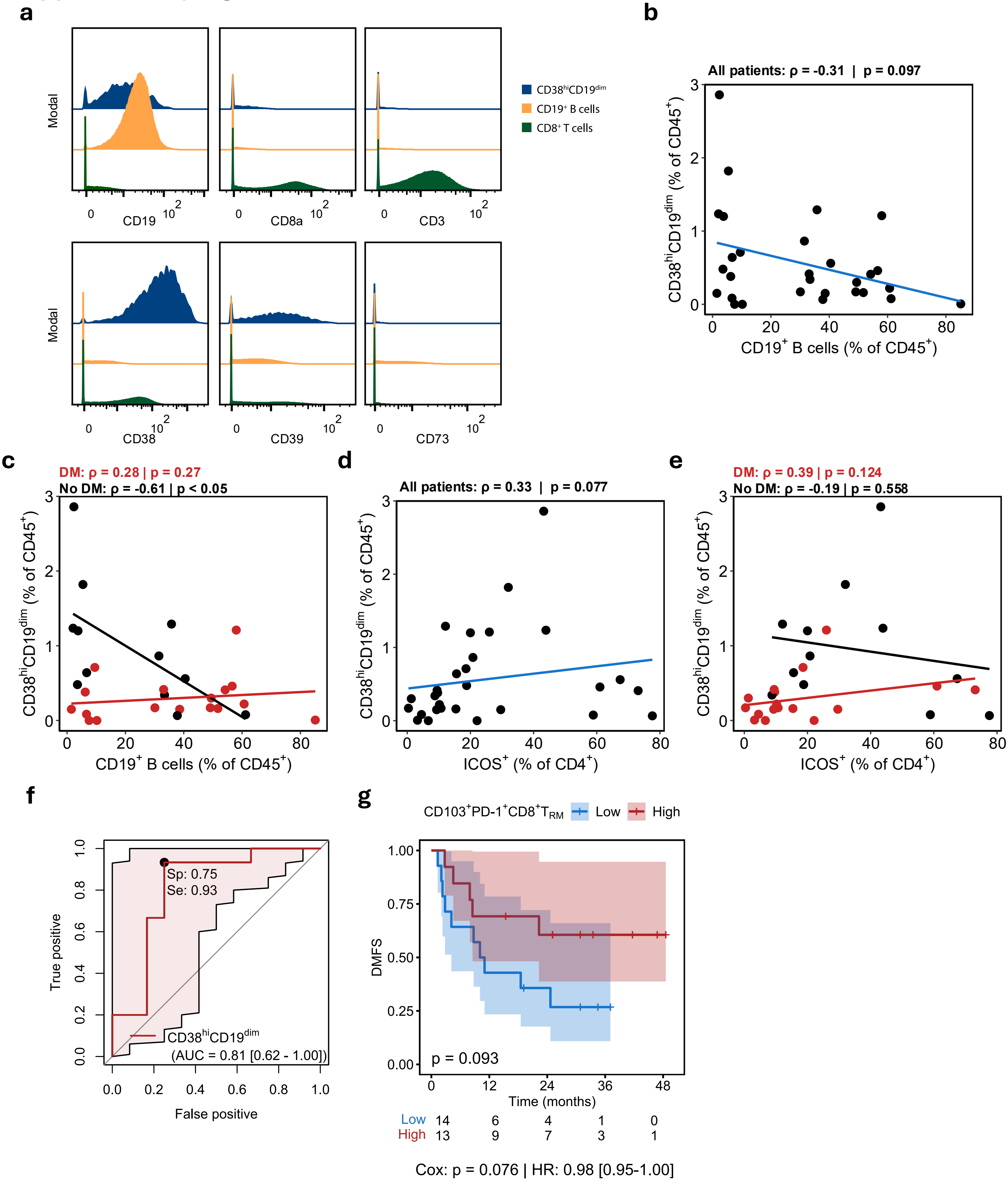


**Supp. Fig. S5**. *$CD38^{hi}CD19^{dim}$ and immune cell correlation.* **a.** Lineage marker expression in the $CD38^{hi}CD19^{dim}$, $CD19^{+}$ B cell and $CD8^{+}$ T cell populations. **b-e**. Spearman correlation of fraction of $CD38^{hi}CD19^{dim}$ and $CD19^{+}$ B cells (**b-c**) or $ICOS^{+}CD4^{+}$ T cells (**d-e**), across all samples (**b, d**) and separated on DM-status (**c, e**). **f.** ROC curve showing the performance of the $CD38^{hi}CD19^{dim}$ population in predicting distant metastasis. True positive (Sensitivity/Se) and False positive (Specificity/Sp) are given on the y- and x-axis, respectively. Area under the curve (AUC) with corresponding 95% confidence interval, Spesifisity (Sp), and Sensitivity (Se) are highlighted. **g.** Kaplan-Meier plot showing distant metastasis-free survival (DMFS) in patients stratified on fraction of $CD103^{+}PD\text{-}1^{+}CD8^{+}$ T-cells higher (red) or lower (blue) than the median. P-value calculated by log-rank test, while p-value for Cox-regression and corresponding Hazard ratio [95% CI] are displayed at the bottom of the plot.

### Supplementary Fig. S6

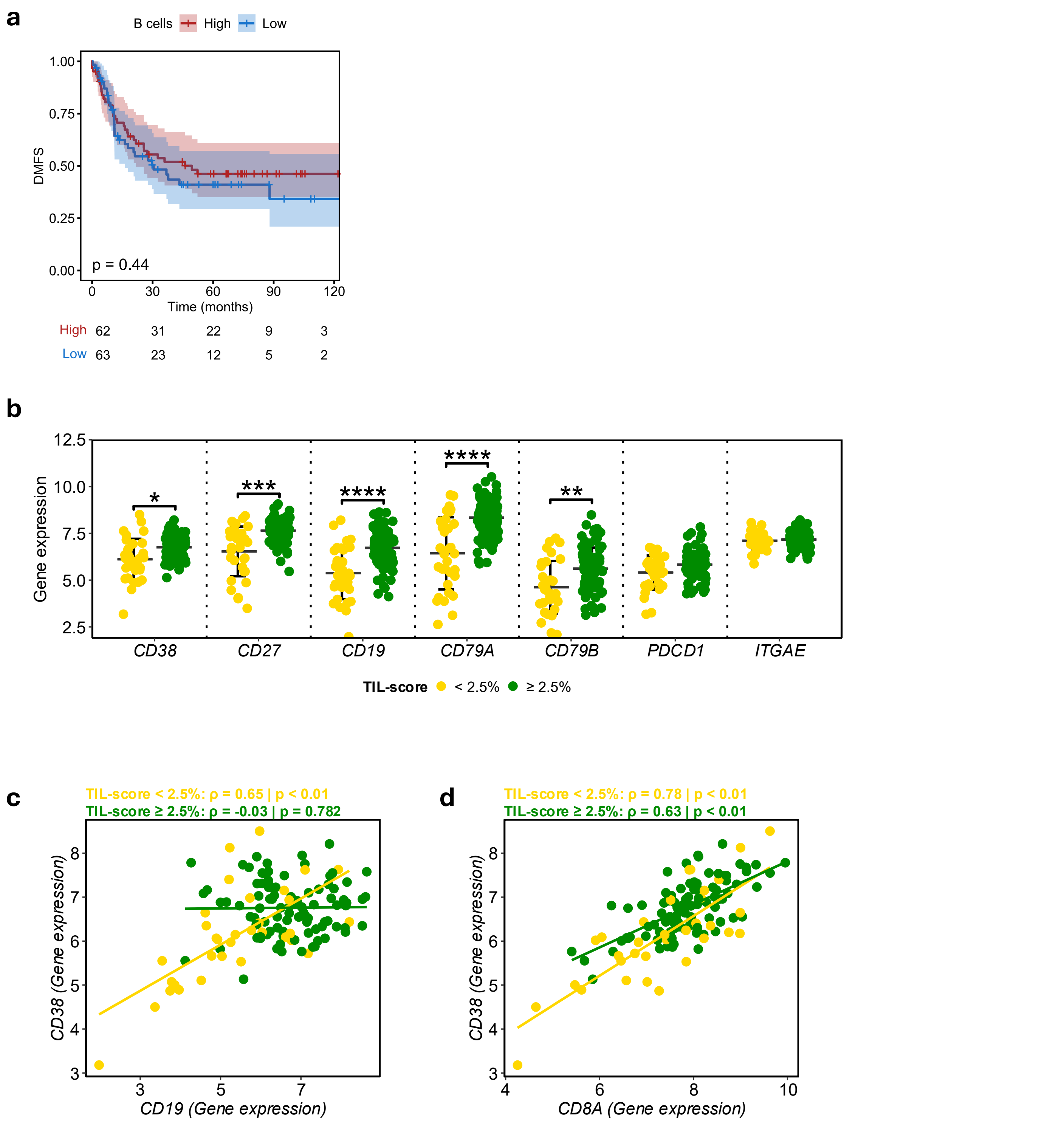


**Supp. Fig. S4**. *B cell survival, and gene expression of B and CD8⁺ T cell markers in untreated samples*. **a.** Kaplan-Meier plot showing distant metastasis-free survival (DMFS) in patients stratified on B cell gene score from NanoString, separated by the median expression into *Low* (< median) and *High* (>median) groups. P-value was calculated by log-rank test. **b.** Expression of selected genes associated with B cell and $CD8^+$ T cell activity in the NanoString cohort, represented as mean±SD. **c-d**. Spearman correlation of *CD38* and *CD19* (**c**) and *CD38* and *CD8A* (**d**) gene expression in the NanoString dataset sepearated into groups defined by TIL-score ≥ 2.5 %.
